# Topographic Organization of Extraoccipital Visual Processing Areas in the Macaque

**DOI:** 10.1101/345363

**Authors:** Gaurav H. Patel, Lawrence H. Snyder, Maurizio Corbetta

## Abstract

The macaque visual system has long been used as a model for investigating the processing of incoming visual stimuli results from the coordinated work of a distributed network of areas interacting at many different levels (Felleman and Van Essen, 1991). While much is known about the organization and layout of the occipital visual areas, there are still substantial gaps in our understanding of layout and organization of the higher-level areas. The goal of this study is to describe the whole brain functional anatomy using BOLD-fMRI in macaques performing a series of demanding visuospatial attention tasks. We wish to study the spatial specificity of visual responses in terms of contralateral preference, i.e. stronger responses to contralateral visual stimuli, as well as retinotopic organization both in terms of polar angle and eccentricity. We found that most visuospatial processing areas only respond to contralaterally presented stimuli; ipsilaterally presented stimuli evoked little or no activity in these areas. Additionally, we found that LIP, MT, and possibly PITd contained polar-angle maps of the contralateral hemifield. These same areas, plus FEF and area 46, appear to have separate representations of the fovea and periphery. When compared to previous human fMRI studies, these results indicate that there may be significant differences between macaque visual processing areas and their putative human homologues.

## Introduction

The processing of incoming visual stimuli results from the coordinated work of a distributed network of areas interacting at many different levels (Felleman and Van Essen, 1991). The macaque visual system has long been used as a model for investigating these processes for two reasons. First, invasive techniques such as single/multi-unit recording and the injection of neuronal tracers allow for detailed investigations of neuronal architecture and function that are not possible in humans. Second, the macaque and human visual systems are thought to be similar in both their psychophysical properties and the organization of the occipital visual areas (Tootell et al., 2003; Orban et al., 2004a; Sereno and Tootell, 2005). In addition to the occipital lobe, visual processing extends into areas within the posterior parietal, inferotemporal, and prefrontal cortices; altogether these areas encompass up to 20% of the macaque cortical surface (Orban et al., 2004a). While much is known about the organization and layout of the occipital visual areas, there are still substantial gaps in our understanding of layout and organization of the higher-level areas.

One of the major limitations of the traditional anatomical and neurophysiological approach is the inability to monitor neural activity simultaneously from the whole brain. This limitation has confounded systematic comparisons of functional and organizational properties of neurons across large cortical distances—within and between areas—because of the differences in the tasks, techniques, and animals used in the studies of each of these areas. As a result, the organization of some of these areas is still disputed, and how this organization changes through the levels of the processing hierarchy is poorly understood. In addition, our knowledge is piecemeal, largely centered on a few selected visual regions (e.g. MT or LIP) that have been object of many investigations, while other areas have been only been cursorily analyzed.

One of the fundamental principles of sensory systems’ organization is that peripheral receptors are mapped onto the brain according to precise rules of topographical organization. For instance, the retina is mapped point-to-point onto the visual cortex, so that the visual cortex represents the visual world in spatial coordinates relative to the center of the retina; this organizing principle is called retinotopy (Wandell et al., 2007). It is well established based on both anatomy and physiology that occipital visual areas are retinotopically organized. Neighboring visual areas can be easily delineated because each area represents one quadrant of the visual world, and the borders between adjacent areas are demarcated by cortex representing a vertical or horizontal meridian. Furthermore, the eccentricity axis of organization, delineating the locations of the foveal and peripheral representations, is orthogonal to the polar angle axis, which defines how the upper, middle, and the lower field representations of the peripheral field are situated relative to one another (Sereno et al., 1995; DeYoe et al., 1996; Wandell et al., 2007).

It is instead less clear if visual areas outside of the occipital lobe are retinotopically organized. Electrophysiology studies have shown that neuronal receptive fields represent increasingly larger portions of the visual field the higher an areas is in the visual hierarchy, and the general agreement is that topography is at best crude in posterior parietal cortex, inferior temporal cortex, and prefrontal cortex (Bruce et al., 1985; Blatt et al., 1990; Boussaoud et al., 1991; Ben Hamed et al., 2001; Sawaguchi and Iba, 2001).

The recent development of monkey functional magnetic resonance imaging (fMRI) recordings of the blood oxygenation level dependent (BOLD) signal provide an indirect measure of neural activity over the whole brain (Logothetis and Wandell, 2004; Viswanathan and Freeman, 2007). Many recent studies have concentrated on the functional anatomy of the occipital lobe, and have used passive visual stimulation to drive visual neurons. However, higher-order visual areas can be typically driven only during active visual tasks, and are much less responsive during anesthesia or to passive stimuli, conditions under which retinotopy has been predominantly described in BOLD-fMRI studies of lower-order visual occipital areas (Brewer et al., 2002; Fize et al., 2003).

The goal of this study is to describe the whole brain functional anatomy using BOLD-fMRI in macaques performing a demanding visual attention task. We wish to study the spatial specificity of visual responses in terms of contralateral preference, i.e. stronger responses to contralateral visual stimuli, as well as retinotopic organization both in terms of polar angle and eccentricity.

## Materials and Methods

### Animal Subjects, Surgery, and Experimental Setup

Two male monkeys (Y and Z, Macaca mulatta, 5-7 kg) were used in accordance with Washington University and NIH guidelines. Prior to training, surgery was performed in aseptic conditions under isofluorane anesthesia to implant a head restraint device. The head restraint was constructed from polyetheretherketone (PEEK) and was anchored to the skull with dental acrylic and 12-16 ceramic screws (4 mm diameter, Thomas Recording GmbH, Germany).

The monkeys were trained to perform the task in a setup that simulated the fMRI scanner environment. In both the training setup and scanner, the animal sat horizontally in a “sphinx” position inside of a cylindrical monkey chair with its head rigidly fixed by the restraint device to a head holder on the chair (Primatrix Inc., Melrose, MA). An LCD projector (BENQ, Irvine, CA) was used to present visual stimuli on a screen that was positioned at the end of the bore, 75 cm from the monkeys’ eyes. A flexible plastic waterspout was positioned near the animal’s mouth for delivery of liquid rewards.

Eye movements were monitored by an infrared tracking system (ISCAN Inc., Melrose, MA). A camera positioned outside of the bore monitored the left eye through a hole in the bottom of the screen; a plastic tube running from the camera to the hole prevented extraneous light from entering the bore via the hole. The eye was illuminated with an IR light source positioned 5-6 cm below the monkey’s left eye. The eye position coordinates were relayed to the behavioral control system, which consisted of two linked computers running custom software. Blinks were detected and the resulting artifacts in the eye-position signal were compensated for on-line.

The monkey was trained to place its paw inside a box that was affixed to the front of the chair under the monkey’s chin, and to rapidly withdraw and replace its paw when a target appeared. These movements were detected by a photoelectric sensor (Banner, Minneapolis, MN). The behavioral control system recorded eye position, hand status and scanner synchronization signals, presented visual stimuli, and delivered rewards. For further details about the experimental setup, see Baker *et al*. (Baker et al., 2006).

### Basic Experimental Design and Training

Each monkey was trained to perform a rapid serial visual presentation (RSVP) target discrimination task. They were then scanned while performing two different versions of the task: one in which a single stream was presented a various locations around the screen, and another in which two streams were presented on either side of the fixation point. Monkey Y was first trained and scanned using the two-stream task, followed by the single-stream task, while monkey Z was first trained and scanned using the single-stream task, followed by the two-stream task. Both animals switched tasks with little or no additional training.

### Single-Stream RSVP Display and Task

At the beginning of a trial, a white fixation point (.3°×.3°) appeared 4.2° vertically below the center of the screen (see **Figure 1.a**). To start the trial, the monkey was given 2 seconds to position his gaze to within 1.3° horizontally and 3.5° vertically of the fixation point, and then another 4 seconds to insert his hand into the hand-response device. The eye-window was longer in the vertical axis to accommodate residual artifacts in the eye-tracking signal stemming from both blinks and pupil constriction. The rapid serial visual presentation (RSVP) stream would then begin coincident with the start of the next MR frames (0-3s).

**Figure 1.**
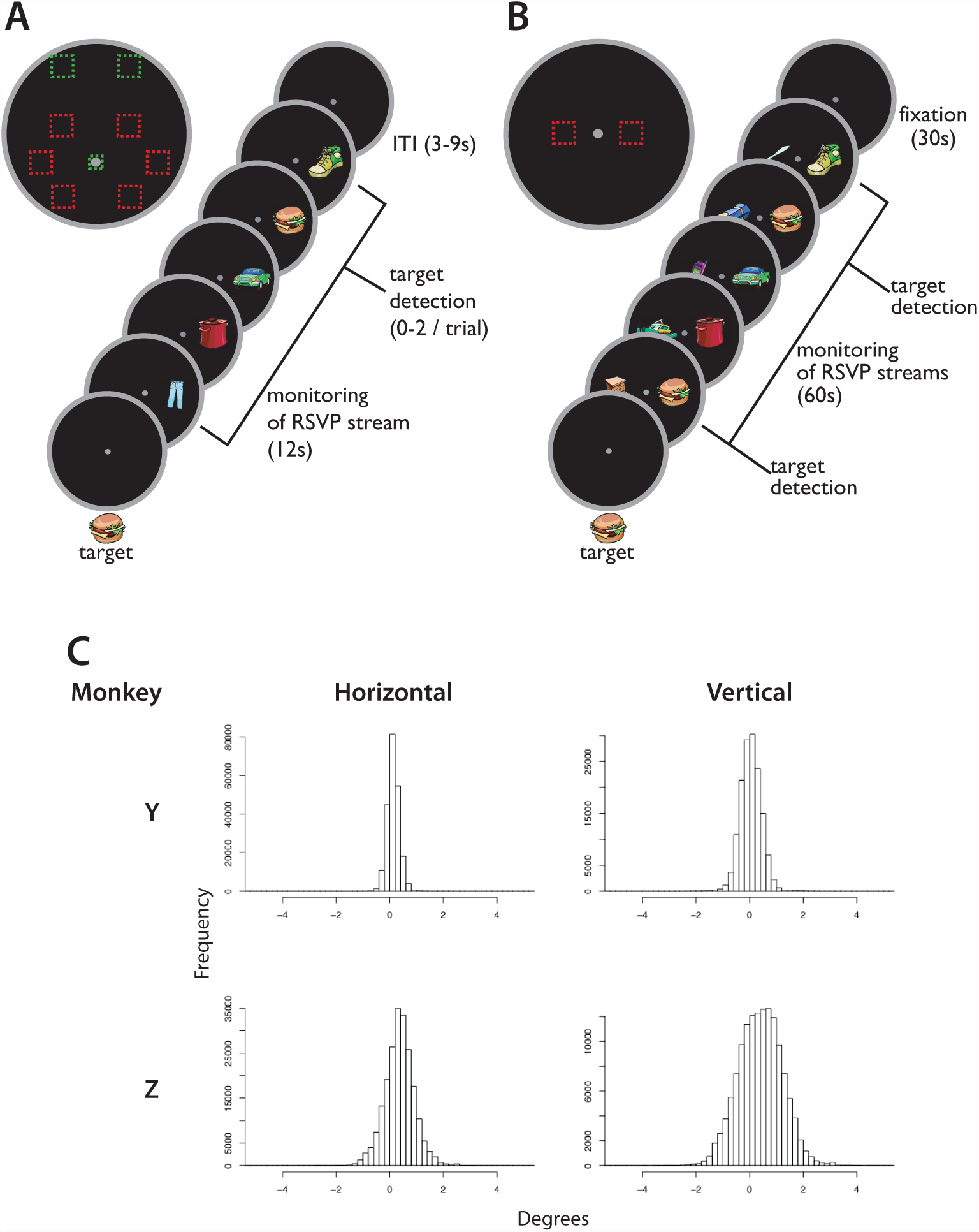
**a)** The single stream paradigm. **b)** The two-stream paradigm. **c)** The distribution of horizontal and vertical eye-positions throughout the single-stream polar angle task, sampled every 40ms. Positive values are in the direction of the presented stream.

The RSVP stream (12 seconds) consisted of bitmap images drawn from a pool of 42 color illustrations of everyday objects (Barry’s Clipart Server, http://www.barrysclipart.com/). For each session, one of the 42 objects was selected a random as the target, and the 41 others were used as distracters. The distracters were shown in random order with the constraint that the same image could not be shown twice in a row. For each BOLD run, we set the time that a single object was on the screen (stimulus duration) as short as possible to keep the monkey’s detection rate between 30 and 100%; between both monkeys the range of durations was 95 to 300 ms.

In the polar-angle version of the single-stream experiment, the stream could appear in one of seven randomly-selected locations: at the fovea (behind the fixation point), or in one of six locations in the periphery in the upper, middle, or lower parts of the left or right visual fields (0°, 45°, 135°, 180°, 225°, or 315° polar angle) centered at 6.8° eccentricity. The size of the objects at the fovea was scaled to 3.6° square, whereas the objects in the periphery were 5.3° square. In the eccentricity version of the paradigm, three locations were used: the fovea; and at 18° polar angle to the left or right of the vertical meridian at 15.6° eccentricity. These positions were chosen as the points on the screen that were as distant as possible from the fixation point. The stimulus size at these peripheral locations was 5.9° square; the visual attributes of the foveal stream were unchanged from the polar-angle version.

The monkey’s task was to detect the memorized target object while maintaining fixation on the central point. A trial could have 0, 1, or 2 targets embedded in the stream with a 25%, 50%, and 25% chance of occurrence, respectively. In a single target trial, the target could appear at any time in the first 10.5 seconds of the 12-second stream. In the two-target trials, the first target would appear within the first four seconds of the start of the stream, and the second 1-6 seconds later. After the appearance of the target, the monkey had 1 second to indicate detection by moving his hand out of and back into the response box. Following each 12-second stream, the monkey was required to maintain fixation for the duration of the inter-trial interval (ITI) of 3-9 seconds.

Upon successful detection, the monkey was rewarded with a drop of juice. For not making any fixation or detection errors during the trial, an additional reward was given at the end of the 12-second stream. Another 1-2 rewards were given for maintaining fixation during the ITI, with the inter-reward interval varying between 0-3 seconds. During some sessions, an additional reward was given at the beginning of the stream. The number and relative sizes of rewards were adjusted to encourage fixation, maximize true hits and minimize false positives, and the size of the rewards was increased with the duration of the session.

If at any time during the trial the monkey broke fixation, or if the monkey removed his hand from the device at a time other than the 1-second reaction time window (such as when falsely detecting a target), the monkey was penalized by blanking the display and aborting the trial. After a short pause, during which no stimuli were presented and no rewards could be obtained, the fixation point reappeared, and a new trial was begun. The duration of the pause varied from 2-9s—as short as possible but long enough to provide enough negative reinforcement to keep the monkey on task. In addition, if the monkey made several errors in a row, the pause was extended to 15-60 seconds. Failures to detect presented targets (misses) were not punished.

At the beginning of the sessions, a sequence of instruction trials lasting ∼20 minutes acquainted the monkey with the session’s target and gave him practice discriminating the target from the distracters. Occasionally the previous session’s target was eliminated as a distracter for the whole session if the monkey kept mistakenly signaling detection for this object during the practice trials.

### Two-Stream RSVP Display and Task

The two-stream task was similar to the single-stream task, except for a few key characteristics and parameters (see **Figure 1.b**). The monkey was required to maintain fixation within 1.1° horizontally and 5° vertically of a central point, but now two RSVP streams were presented simultaneously: one at 5.3° eccentricity horizontally to the left of the fixation point, and the other at 5.3° to the right. The streams in this experiment were moved closer to the fixation point than in the single-stream task because monkey Y initially had difficulty performing the task with the streams at 6.8° eccentricity (however, these fixation breaks were relatively rare; see **Results**). The sizes of the objects were scaled to 4.6° square. Once the streams started, they remained on-screen for 60 seconds. In the streams, targets appeared once every 6 +/− 3 seconds, and the stimulus duration of the objects was varied between 45 and 225ms; the duration was set during the warm-up trials so that the monkey was detecting 50-80% of the presented targets. Each day, one of the streams was designated as the “target” stream, and all of the targets in that day appeared in that stream. The “target” stream was chosen randomly each day, and the monkey learned which was the designated side during the warm-up trials. Besides the appearance of the targets, the two streams did not differ in any way in visual stimulus parameters. The reinforcement schedule was also similar, except that the bonus rewards were given after every 2-3 target presentations. After 60 seconds, the streams remained off for a 30 second fixation period, during which the monkey was given a reward every 6 +/− 3 seconds for maintaining fixation. In monkey Z, an additional fixation reward was given every 6 seconds. Then the cycle would repeat until either the speed needed to be adjusted or the monkey was finished for the day.

### Data Collection

Functional and anatomical data were collected in a Siemens 3T Allegra MRI scanner (Siemens Medical Solutions, Erlangen, Germany). High-resolution structural images were collected in separate sessions, during which the animal was chemically restrained (10 mg/kg ketamine, 0.6 mg/kg xylazine, .05 mg/kg atropine). T1-weighted images were acquired using a magnetization-prepared rapid acquisition gradient-echo pulse sequence [MP-RAGE; (0.5 mm)^3^ isotropic resolution, flip angle = 7°, six acquisitions] and a volumetric transmit and receive coil (16 cm i.d.; Primatrix).

Functional data were collected using a gradient-echo echo-planar pulse sequence sensitive to BOLD contrast (T2*) (T2* evolution time = 25 ms, flip angle = 90°) and a transmit-receive surface coil (13 cm inner diameter; Primatrix). The coil fit around each animal’s head post and was saddle-shaped to provide more extensive brain coverage as compared to a planar surface coil. Fifty-two coronal slices, each with a square field of view (96 × 96 mm, 64 × 64 base resolution, dorsal-to-ventral phase-encoding) and a thickness of 1.5 mm, were obtained using contiguous, interleaved acquisition, and a volume repetition time (TR) of 3000 ms. This scanning protocol was chosen to cover the whole brain at an isotropic spatial resolution of (1.5 mm)^3^. The first four volumes of each run were excluded from the analyses to allow for the equilibration of longitudinal magnetization. A T2-weighted image was acquired at the beginning of one session using a turbo spin-echo sequence [TSE; (1×1×1.5mm)^3^ resolution, flip-angle = 150°, single acquisition).

In the single-stream polar angle experiment, up to 200 volumes were acquired in each fMRI run; runs were cut short if the monkey stopped performing the task. A total of 7500 frames in 39 runs across 7 sessions were collected from monkey Y, and 6700 frames in 39 runs across 18 sessions from monkey Z. This resulted in about 100 successfully completed trials collected for each of the seven stream positions in each monkey. In the eccentricity experiment, 23 runs (4400 frames, 100 trials per position) were collected in monkey Y and 30 runs (5900 frames, 200 trials per position) in monkey Z.

In the two-stream experiment, up to 1409 frames (71 minutes) were collected in a single run; the scanner was only stopped if the stimulus parameters needed adjustment or if the monkey quit working. In monkey Y, a total of 5800 frames were collected in 11 runs in 9 scanning sessions, and in monkey Z a total of 7100 frames were collected in 8 runs in 8 scanning sessions.

### Preprocessing

In the polar angle and eccentricity experiments, all runs in which the monkey successfully completed more than 50% of the presented trials were included in the analysis (37/39 runs in monkey Y, 35/39 runs in monkey Z). The exception was monkey Y’s eccentricity data; this criterion was removed due to his tendency to increased errors resulting from quick out-and-back saccades to the peripheral streams. Because the error trials were modeled separately, the removal of this criterion did not appear to affect the data quality, as the statistical, focality of activations, and overall distribution of activity appeared to be similar in the two monkeys in this experiment. No runs were excluded for excessive movement. In the two-stream experiment no runs/sessions were excluded due to excessive movement or poor performance.

Each reconstructed fMRI run produced a 4-dimensional (x, y, z, time) data set that was passed through a sequence of unsupervised processing steps using in-house software. The data were first corrected for asynchronous slice acquisition using cubic spline interpolation, and also for odd-even slice intensity differences resulting from the interleaved acquisition of slices. Correction factors were then calculated for 1) normalization across runs, in which each four-dimensional data set was uniformly scaled to a whole brain mode value of 1000; and 2) a 6-parameter rigid body realignment to correct for within- and across-run movement. Finally, correction factors were calculated for aligning the data from each session first to each other and second to the monkey’s own T2-weighted image, aligned to the macaque F6 atlas (http://sumsdb.wustl.edu/sums/archivelist.do?archive_id=6636170) using a 12-parameter affine registration algorithm (Snyder, 1995). All of these correction factors were then applied to the data in a single resampling step. To further refine the cross-session alignment within each monkey, an additional recursive alignment algorithm was implemented. Atlas-aligned representative images from each session were averaged together, and then each session’s representative image was realigned to this average using the 12-parameter affine registration. These newly aligned images were then averaged together and realigned to the T2-weighted image, which in turn was used as the target for the next realignment iteration. This was repeated until the change in variance averaged across all the voxels across all of the sessions asymptotically reached a minimum. Finally, the final atlas-registration matrix and the above-calculated normalization and movement-corrections factors were re-applied to the data in a single resampling step.

### Projection of the statistical data to the cortical surface

For each monkey, a cortical surface model was created by segmenting the gray and white matter of the monkey’s own MPRAGE (http://brainvis.wustl.edu, (Van Essen, 2002)). The resulting segmentation volume was then edited to fit the average atlas-aligned EPI image that resulted from the above-described atlas alignment scheme. The cortical surface was then flattened and registered to the macaqueF6 surface atlas (http://sumsdb.wustl.edu/sums/archivelist.do?archive_id=6636170) using sulcal-folding markers. To project statistical maps (see below) to the flattened, registered cortical surface, the volume maps were resampled to (.5mm)^3^, and then each point on the surface was painted with the highest absolute z-score within 1.5 mm. For comparisons between the four hemispheres, these maps were then warped to the macaqueF6 atlas right hemisphere surface by way of the sulcal-folding surface registration procedure described above.

### Analysis of Contralateral Preference

Statistical maps of activity were created using a general linear model implemented with in-house software. To create z-statistic maps, each event-type was modeled as an independent regressor formed by convolving a gamma function with a 2-seond delay (Boynton et al., 1996) with a boxcar of a specified duration. In the single-stream experiment, the included regressors were the 12-second stream for each location, target detections (.5 seconds), bonus rewards (.5 seconds), and missed targets (.5 seconds) (see **Figure 1.a**). In addition, RSVP streams cut short by errors were coded separately from completed (12-second) streams, and regressors were included for the fixation break and false-detection error pauses. If the punish period lasted longer than 21 seconds, those frames were excluded from the analysis. In the two-stream experiment, reward and punish events were similarly modeled (see **Figure 1.b**). The streams were separated into ones that were 60 +/− 3 seconds in duration and ones that were less than 60 seconds due to errors. In addition, the streams were coded separately by whether the session was an “target-left” or “target-right” session. In both models, linear trend and baseline terms were also included as regressors. In the model, only the amplitude of the regressors was allowed to vary freely for each voxel, and the standard error was estimated from the remaining variance. From these amplitudes and standard errors, a z-score of each regressor’s fit to the voxel’s time-course was estimated. Contrasts between responses to event-types were calculated as the z-score of the difference between the magnitudes relative to the fixation baseline. To estimate the time-course of the response to a given event, separate models were created in which no response was assumed for the fixed-length event-types.

While the statistical maps from the two experiments were strong, each map also contained a number of extraneous foci that were not consistent between experiments and in some cases appeared to be artifactual. In order to limit the effect of this noise on the further analyses, we created a mask for each monkey consisting of voxels that were a) significantly activated in the two-stream experiment by the presence of the streams versus the fixation baseline and b) significantly activated by the streams presented in one half of the visual field over the other in the single-stream polar angle experiment. The latter voxels were isolated by summing the contrast maps generated by the three left visual-field stream regressors, subtracting the sum of the three right visual-field stream contrast maps, and taking the absolute value of the result. The two maps that resulted from (a) and (b) were then each thresholded at p<.05 (Bonferroni correction: z>4.7). Finally, the logical conjunction of the two thresholded maps (‘and’ operation) was used to mask the fixed-effects volume map from each of the two monkeys.

This process resulted in a map contained statistically significant BOLD signals that were similar in distribution in all four hemispheres. For additional analyses, regions of interest (ROIs) were created from this map in each monkey using an automated peak-search algorithm that grouped together voxels within 6mm of a local maxima within the conjunction map. This procedure, however, sometimes arbitrarily divided clearly contiguous foci of activity. To remedy this, the various atlases available on the macaqueF6 surface in Caret and the Saleem and Logothetis atlases were used to combine these automated ROIs. In general, unless clear anatomical criteria could be used to divide neighboring ROIs into separate clusters, adjacent clusters were combined to form a single ROI. These combined ROIs were then named according to the atlases. This procedure gave us four occipital (V1, V2/V3d, V2/V3v, V4) and 7 extra-occipital (LIP, MT, PITd, CITd, AITd, FEF, and area 46p) ROIs for each of the four hemispheres. The only exceptions were a lack of a V1 ROI for monkey Z’s left hemisphere and a CITd in monkey Z’s right hemisphere. For each ROI, we extracted the average time-course of the response to the RSVP stream at each location in the polar angle experiment. Time-points 2-5 were averaged together for each location to create an estimate of the BOLD-response magnitude. Then, for each ROI, these magnitudes were averaged together for the three contralateral and also for the three ipsilateral locations. These magnitudes were used to create an index of the preference for the contralateral visual field over the ipsilateral field ((contra + ipsi)/contra). These indices were then averaged for each ROI across the 4 hemispheres.

### Analysis of Polar Angle Maps

To cleanly exhibit any retinotopic maps on the cortical surface, a new GLM was created in which the monkey’s own BOLD response was used as the assumed hemodynamic response function. The assumed response was the average time-course of the BOLD response to the three contralateral field stream locations from LIP, MT, PITd, CITd, AITd, FEF, and area 46p; the regressors in the model were otherwise identical to the GLM described above. This model gave more accurate estimates of the actual BOLD responses in each monkey to the contralateral-field RSVP streams than the model with standard assumed responses, but still did not bias the resulting maps toward any single contralateral visual field location. This GLM was then used to perform two analyses, whose combined results were used to identify retinotopic maps.

The first analysis consisted of contrasting the activity evoked by each stream location with the average of the three ipsilateral field locations. This contrast resulted in less-noisy statistical maps than contrasting the stream activity with the fixation baseline, as the contra-ipsi contrast removed activations common to the presentation of streams irrespective of visual field location. The six resulting statistical maps and the corresponding magnitude maps were projected to the macaqueF6 surface (see above). A fixed-effects map was created from the four hemispheres for each of the three contralateral stream location. These three maps per hemisphere were simultaneously projected to the cortical surface, and the statistical threshold was raised until the peaks and surrounding surface tiles were isolated for each stream location (theshold was at least p<.05 Bonferroni correction). A border was then drawn around these tiles for each location representation to create surface ROIs representing each visual field location. For each ROI, the average magnitude of activation evoked by each stream location was extracted and averaged across hemispheres.

The second analysis consisted of contrasting the activity evoked by the upper and lower streams in the contralateral field for each hemisphere. These z-statistic maps were then projected to the macaqueF6 surface, and then the data from the four hemispheres were combined to create a fixed-effects map. A threshold of p <.05 (Bonferroni corrected) was used to isolate the maxima (upper field) and minima (lower field). Since the first analysis was only qualitative, this analysis was used to verify that the maps seen in the first analysis were statistically reliable.

### Analysis of the Periphery vs. the Fovea

To search the cortical surface for separate foveal and peripheral representations within a single area, a GLM was used to create z-statistic contrast maps of the activity evoked by the peripheral and foveal streams in the eccentricity experiment. Since there was no unbiased method for using the monkeys’ own responses in the GLM, a standard gamma-function HRF was used (Boynton et al., 1996). The regressors used for each event type in the model were otherwise identical to those used in the polar-angle experiment. For each monkey, the two resulting contrast maps as well as the corresponding magnitude maps for each stream location were projected to the macaqueF6 surface. The four contrast surface maps were combined to create a fixed-effects map, and then thresholded to reveal the statistically significant differences in the contrast (p<.05, Bonferroni correction). Borders defining surface ROIs were then drawn around the positive (peripheral) and negative (foveal) clusters that fell in or near each contralateral-preferring ROI. To determine the strength of the representations for each ROI, the difference in the magnitude of activity evoked by the contralateral peripheral and foveal streams was averaged across the four hemispheres for both the contralateral peripheral and foveal locations.

### Eye position analysis

Although large eye position windows were enforced during data collection, actual eye position was generally very close to the fixation point. In order to quantify this, we plotted histograms of the horizontal and vertical components of the monkey’s eye-position calculated in 40ms bins, separately for the one- and two-stream experiments. The components were plotted as the horizontal and vertical displacement towards the presented stream; for the horizontal meridian streams, the vertical components were not included. The resulting eye-position distributions were used to calculate the mean and standard deviation of the mean horizontal and vertical eye-position for each monkey in the polar angle and two-stream experiments.

## Results

To study the topographic properties of macaque cortical visual processing areas, we trained two macaque monkeys to perform two versions of a visual search task. In the first experiment, the monkey fixated a central point while a single RSVP stream appeared for 12 seconds at the fovea or in the periphery (see **Figure 1.a**). The peripheral streams appeared at 6.8° eccentricity in one of six locations in the periphery in the upper, middle, or lower part of either the left or right visual field. In the second experiment, two streams were shown simultaneously for 60 seconds, one on each side of the fixation point (see **Figure 1.b**). In each experiment, the monkey’s task was to detect the target image, which was chosen randomly from the 42 possible images each session.

### Behavior

Across the three experiments, the mean detection rates and reaction times were 49% and 515ms for monkey Y, and 76% and 470ms for monkey Z (see **Table 1** for further details). With only one exception, each monkey successfully completed (did not break fixation or make a false detection) >50% of the presented trials in all three experiments. (The one exception was that monkey Y successfully completed only 43% of the eccentricity trials). In the analysis of the BOLD data, the signals related to errors were explicitly modeled and removed from the signals of interest.

**Figure 1.c** shows horizontal and vertical fixation error in both monkeys while performing the polar angle experiment (40ms bins). Horizontal fixation was good (monkey Y: mean fixation error of .13° (sd=.26°); monkey Z: .36° (58°)), but the vertical error in monkey Z (.39° (.92°) was much larger than in monkey Y (.03° (.52°)). In both monkeys, the tolerance limit for vertical errors was greater than the horizontal because of eye-tracking artifacts. However, only monkey Z appeared to use this to his advantage, and that was largely for the upper-right and upper-left stream positions. However, even the largest errors were less in eccentricity than the inner edge of the peripheral streams (4.15° eccentricity). In the two-stream experiment, monkey Y and Z’s gaze deviated .16° (.26°) and .09° (.36°), respectively, toward the attended stream, indicating tight control of fixation.

**Table 2.1.**
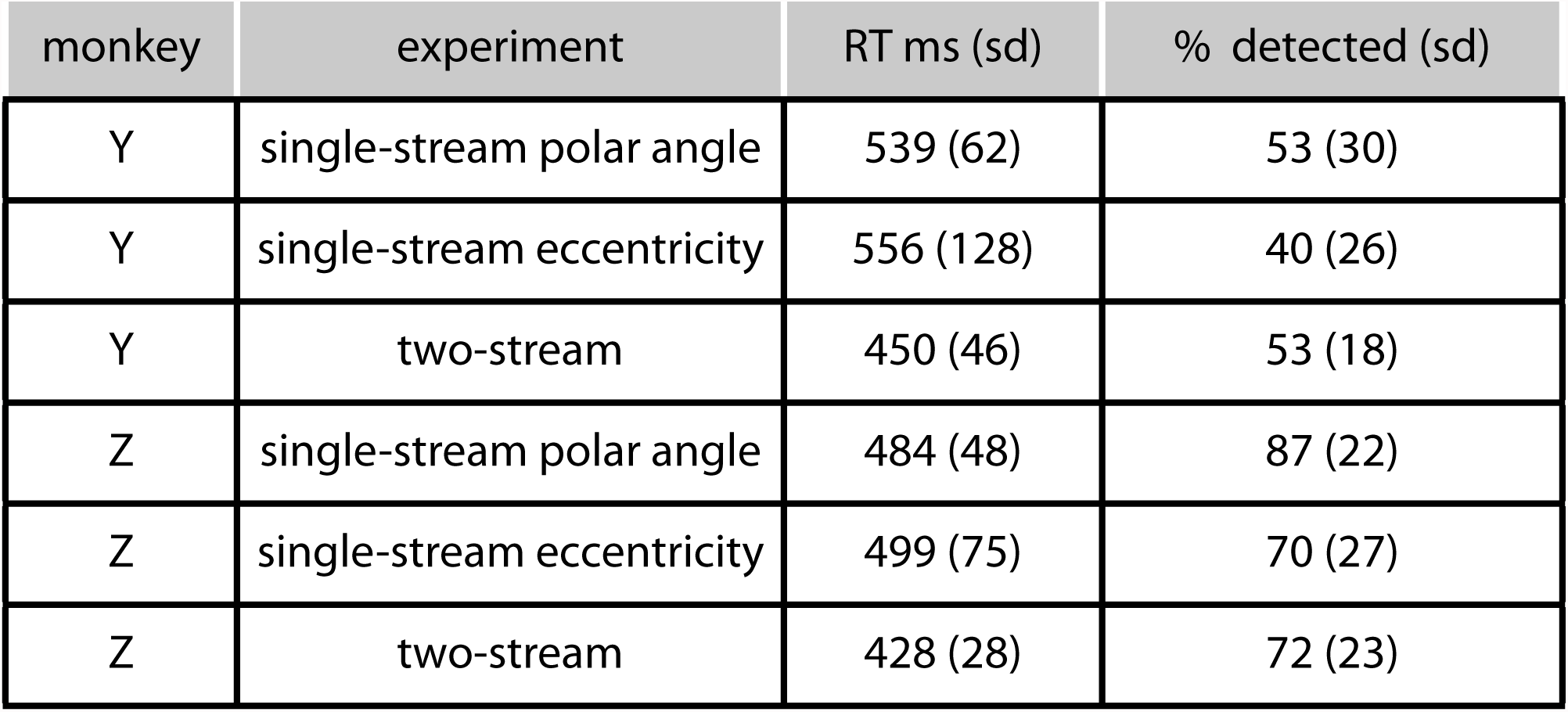
Target detection rates and reaction times for both monkeys in the three experiments.

| monkey | experiment | RT ms (sd) | % detected (sd) |
| --- | --- | --- | --- |
| Y | single-stream polar angle | 539 (62) | 53 (30) |
| Y | single-stream eccentricity | 556 (128) | 40 (26) |
| Y | two-stream | 450 (46) | 53 (18) |
| Z | single-stream polar angle | 484 (48) | 87 (22) |
| Z | single-stream eccentricity | 499 (75) | 70 (27) |
| Z | two-stream | 428 (28) | 72 (23) |

### Location of Evoked Activity

We used a GLM to compare the BOLD response to contralateral versus ipsilateral streams in the single-stream experiment, and to model the combined response to paired streams in the two-stream experiment. In the resulting z-statistic maps, we observed foci of BOLD signals in occipital, posterior parietal, inferotemporal, and prefrontal cortices in both monkeys (see **Figures 2.a** and **2.b** for the fixed-effects surface maps representing all four hemispheres for each condition, p<.05 Bonferroni correction). The conjunction of these maps from the two experiments after multiple-comparisons correction (**Figure 2.c**) was used for further analyses to simplify region of interest (ROI) selection. **Figures 2.d** and **2.e** (volume conjunction maps from each animal) demonstrate that these ROIs were remarkably consistent between the four hemispheres in the two monkeys.

**Figure 2.**
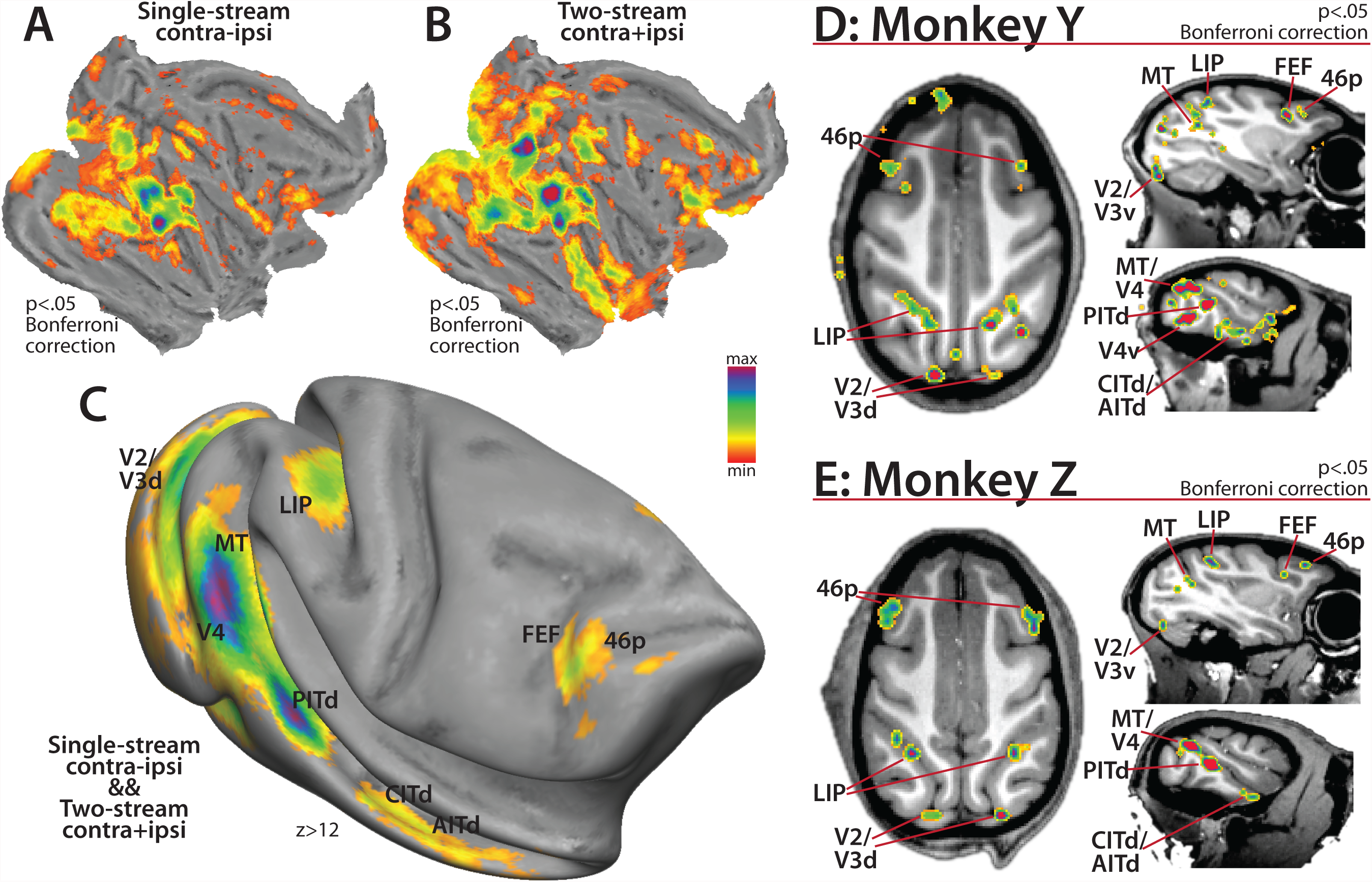
Distribution of cortical activity in the single-stream polar angle and two-stream experiments. **a)** & **b)** Fixed effects average of activity evoked by the single- and two-stream experiments projected to the flattened macaqueF6 surface. **c)** Fixed effects average of the conjunction of the voxels significantly activated in the contrasts in a) and b) projected to the inflated macaqueF6 surface. **d)** & **e)** Horizontal and saggital sections showing the voxels demonstrating significant effects for both the single-stream and two-stream contrasts in each individual monkey.

To verify that the appropriate parts of the retinotopic map were stimulated by the two paradigms, we first examined the occipital foci in relation to previously documented anatomical landmarks/retinoptopic maps correspondences. Foci in the conjunction maps in all four hemispheres were observed in the caudal portion of the calcarine sulcus, the posterior bank of the lunate sulcus, in the fundus of the inferior occipital sulcus, and on the gyral surface between the lunate and superior temporal sulci. These corresponded in location to the horizontal meridian representations in V1, between V2d and V3d, between V2v and V3v/VP, and in V4, respectively. In monkey Y, an additional focus was seen at the ventral anterior border of V4 (labeled V4v in **Figure 2.d**) but did not appear in monkey Z; this discrepancy was most likely due to differences in the susceptibility artifact stemming from the nearby ear canals. In the single-stream paradigm, the upper field streams evoked activity in ventral visual cortex locations between the horizontal meridian representations, and the lower streams in their corresponding locations in dorsal visual cortex. After the reliability of the activations had been verified in occipital cortex, the loci of activity in the rest of cortical surface were compared to the various partioning schemes available on the macaqueF6 surface and also the Saleem and Logothetis atlas (Van Essen, 2002; Saleem and Logothetis, 2006).

In the posterior parietal cortex, a focus of activity in the conjunction map spanned much of the lateral bank of the IPS, from the fundus to the gyrus, and stretched ∼1 cm in the axis parallel to the fundus starting at the bend of the IPS. This focus covered much of the areas LIPd and LIPv in the Lewis and Van Essen atlas. In monkey Y, another focus of activity was observed on the lateral bank at the junction of the parietal-occipital and intraparietal sulci, but the presence of this focus was inconsistent in monkey Z, and was not considered further.

In the superior temporal sulcus, the most caudal (posterior) focus of activity was immediately rostral (anterior) to V4. The peak voxels fell near the border of V4tp/a and MT in the Lewis and Van Essen atlas; after consulting the Saleem and Logothetis atlas we labeled the ROI MT. This focus stretched about 7mm along the lateral-inferior bank of the caudal portion of the STS. Another 7mm rostral to MT was another large focus of activity whose boundaries fit within PITd in the Felleman and Van Essen atlas. PITd was about 5mm long in the rostral-caudal axis, and on the lateral portion of the inferior bank of the STS, near the lip of the gyrus. 5mm rostral to PITd in 3 of the 4 hemispheres was another pair of foci of activity (in monkey Z’s right hemisphere there was only one focus 10mm rostral to PITd). These two regions also fit well with Felleman and Van Essen areas CITd and AITd. Both of these foci also fell along the lateral (superficial) portion of the inferior bank of the STS, and extended rostrally nearly to the temporal pole.

In prefrontal cortex, two main foci of activity were observed in all four hemispheres. The first was along the anterior bank of the arcuate sulcus, stretching from the midpoint into the inferior ramus 4mm laterally. This focus was split between the 8Ac and 6Vam in the Lewis and Van Essen atlas and corresponded to FEF in the Saleem and Logothetis atlas. Anterior to FEF was a focus that extended across the gyrus anterior to the arcuate sulcus and into the posterior portion of the principal sulcus. This focus corresponded to the ventral portion of area 46p in the Lewis and Van Essen atlas. Additional foci were observed in both experiments near the inferior termination of the arcuate sulcus in all four hemispheres, but the results from the two experiments did not align well enough in monkey Z to survive the conjunction, and this area was not considered further.

The conjunction maps in each monkey (**Figures 2.d** and **2.e**) were used to create volume-space ROIs for each monkey to extract BOLD magnitudes and time-courses from the polar angle experiment for the analysis of contralateral bias. The fixed-effects conjunction map on the macaqueF6 surface (**Figure 2c**) was used to draw borders for comparison with the retinotopic maps derived from both the polar angle and eccentricity experiments discussed below.

### Contralateral Preference

For each of the volume-space ROIs in each hemisphere, we extracted a time-course of the BOLD modulation evoked by the presentation of the RSVP stream for each of the six peripheral stream positions in the polar angle experiment (see **Figure 3.a** for examples from PITd from all four hemispheres). These time-courses demonstrate that the contralateral streams evoke a sustained 12-second BOLD response that is remarkably consistent across the three stream positions (solid lines in **Figure 3.a**), while the responses for the three ipsilateral streams are relatively flat (dashed lines in **Figure 3.a**). For each monkey, the basic shape of the time-courses was consistent across all of the ROIs. The time-courses were used to compute a laterality index, defined as the difference in the mean magnitude of the BOLD signal evoked by contralaterally-versus ipsilaterally-presented streams, normalized by the mean contralateral stream magnitude. An index value near zero meant that the magnitude of the response in the ROI was equivalent for stimuli presented in the contralateral and ipsilateral fields, whereas an index value near 1 meant that the response was strongly biased towards the contralateral field with almost no response to ipsilaterally-presented stimuli. The contralateral index values were averaged across hemispheres for each ROI (see **Figure 3.b**). In general, all of the ROIs, both occipital and extra-occipital, had index values near or above 1 (and not significantly different from one), meaning these areas were not activated by ipsilaterally-presented streams. The exceptions were CITd and AITd; the trend in the three inferotemporal ROIs indicated declining contralateral preference with increasing rostral location. The large negative value and standard error for AITd is a result of a small contralateral response in one hemisphere, so that normalization resulted in a very large negative index. Since the ROI selection was biased towards voxels with a contralateral preference (see **Methods**), we repeated the index calculation for the unbiased ROIs selected from the map of voxels significantly activated in the two-stream paradigm alone. While the results are more variable between hemispheres, the mean index values for the extra-occipital ROIs are still near 1, with the same trend in inferotemporal cortex as above (see **Figure 4**).

**Figure 3.**
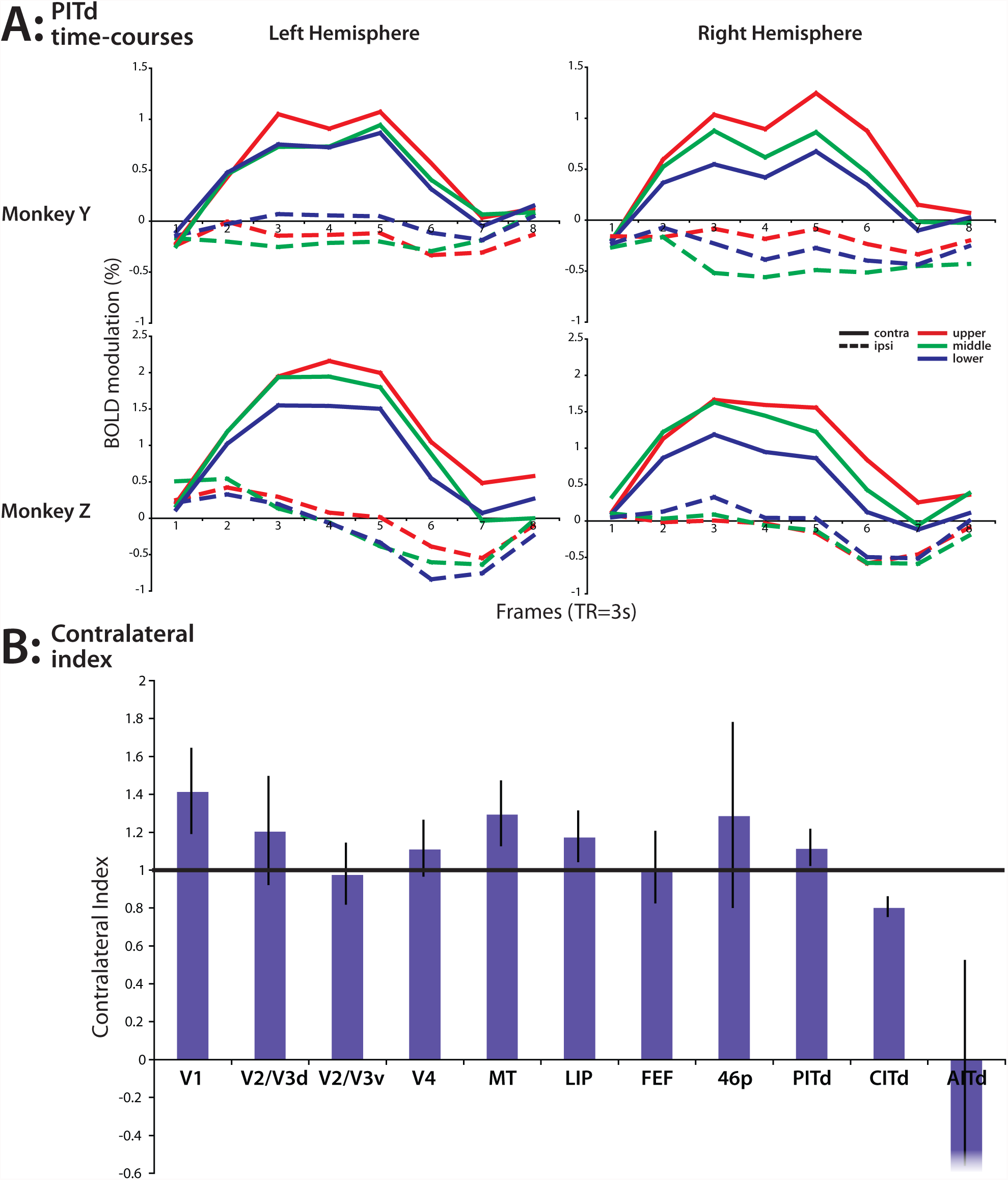
Contralateral bias in the single-stream polar angle experiment. **a)** Time-courses of the BOLD signal in PITd in the four hemispheres evoked by the peripheral RSVP streams. **b)** The contralateral bias of all of the visual processing areas (index = (contra-ipsi)/contra).

**Figure 4.**
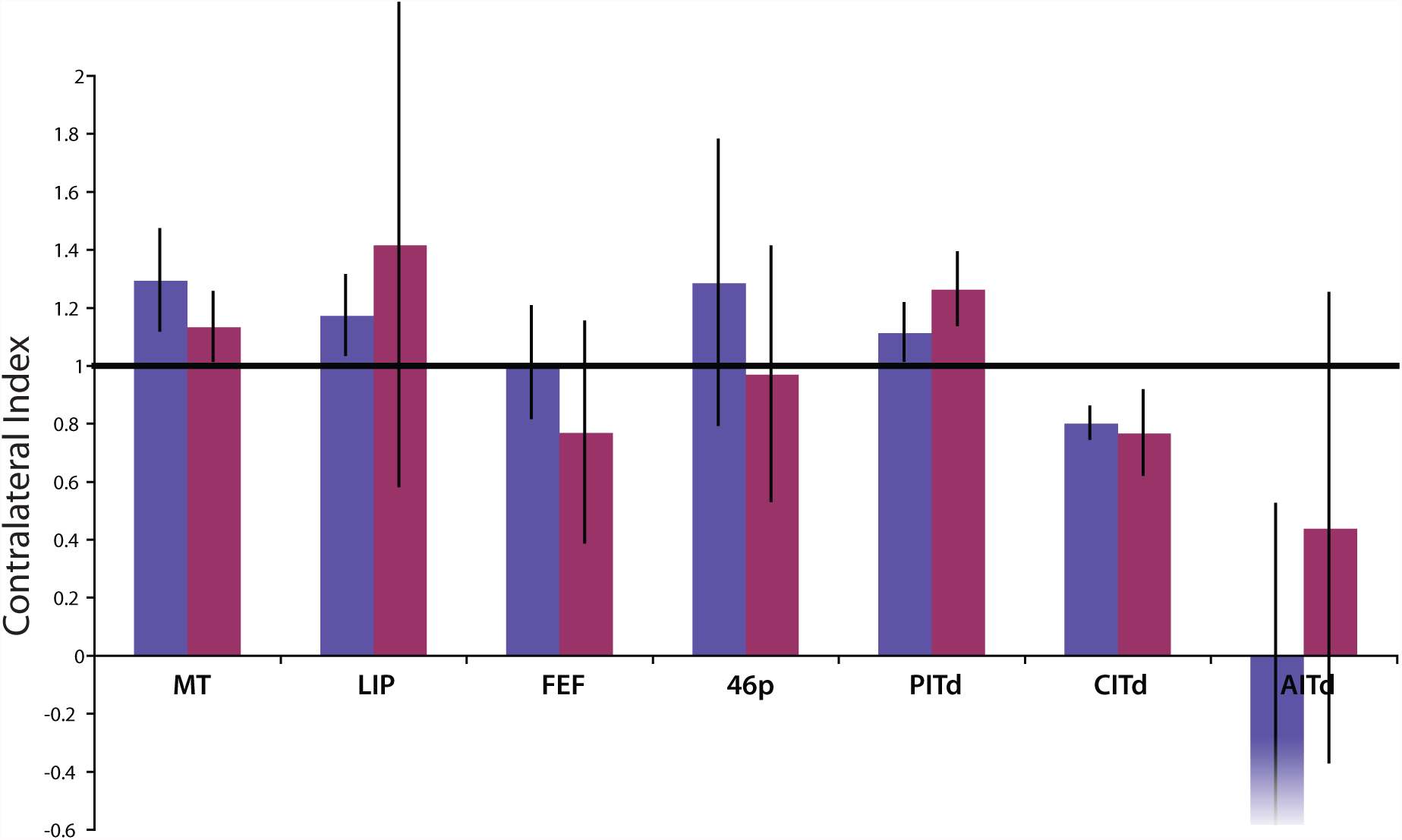
Comparison of the contralateral bias of extraoccipital areas in ROIs defined by the conjunction of the single-stream C-I and two-stream C+I contrasts (blue-purple) and by the two-stream C+I contrast alone.

### Polar Angle Map

To determine which areas contained retinotopic maps of the contralateral field, we performed two separate analyses. The first analysis isolated the activity specific to each of the three contralateral streams by contrasting the activity evoked by each contralateral stream with the average activity evoked by the three ipsilateral streams; these maps were used to determine if the maps were contiguous representations of the contralateral hemifield (p<.01 uncorrected). The second analysis was a voxel-wise contrast directly comparing the activity evoked by the contralateral upper- and lower-field streams for each hemisphere, and was used to determine the statistical strength of the maps (p<.05, Bonferroni correction). The projections of these data to the macaqueF6 surface were used to search for statistically reliable maps within the boundaries of the conjunction ROIs, and to create fixed-effects maps of each contrast.

We found three areas containing retinotopic maps of the contralateral field that reproduced reliably in all four hemispheres: LIP, MT, and PITd. In LIP, the activity evoked by the upper visual field stream was caudal to that of the lower visual field stream, and the peak of the middle visual field stream activity fell between the upper and lower field peaks in all four hemispheres (Patel et al., 2010 Figure 3a). The fixed-effects maps underscored the presence of a contiguous map of the contralateral peripheral visual field with an axis of organization paralleling the fundus (Patel et al., 2010 Figure 3b) In MT, the axis of organization ran mostly rostral-caudal, with the upper field representation rostral and slightly medial to the lower and middle field representations (see **Figures 5.a** and **5.b**). The middle field representation often appeared to be closer to V4 than the upper or lower field representations, and in some hemispheres this appeared to be a shared horizontal meridian representation with V4. In PITd, the map in all four hemispheres also ran rostral-caudal, with the lower field representation at the rostral end (see **Figures 6.a** and **6.b**). However, the upper field representations required a higher threshold to isolate the peak than the other representations, implying that it had more power than either the middle or lower field representations. We did not find evidence of any other contiguous maps of the contralateral field in the other ROIs or elsewhere in extra-occipital cortex.

**Figure 5.**
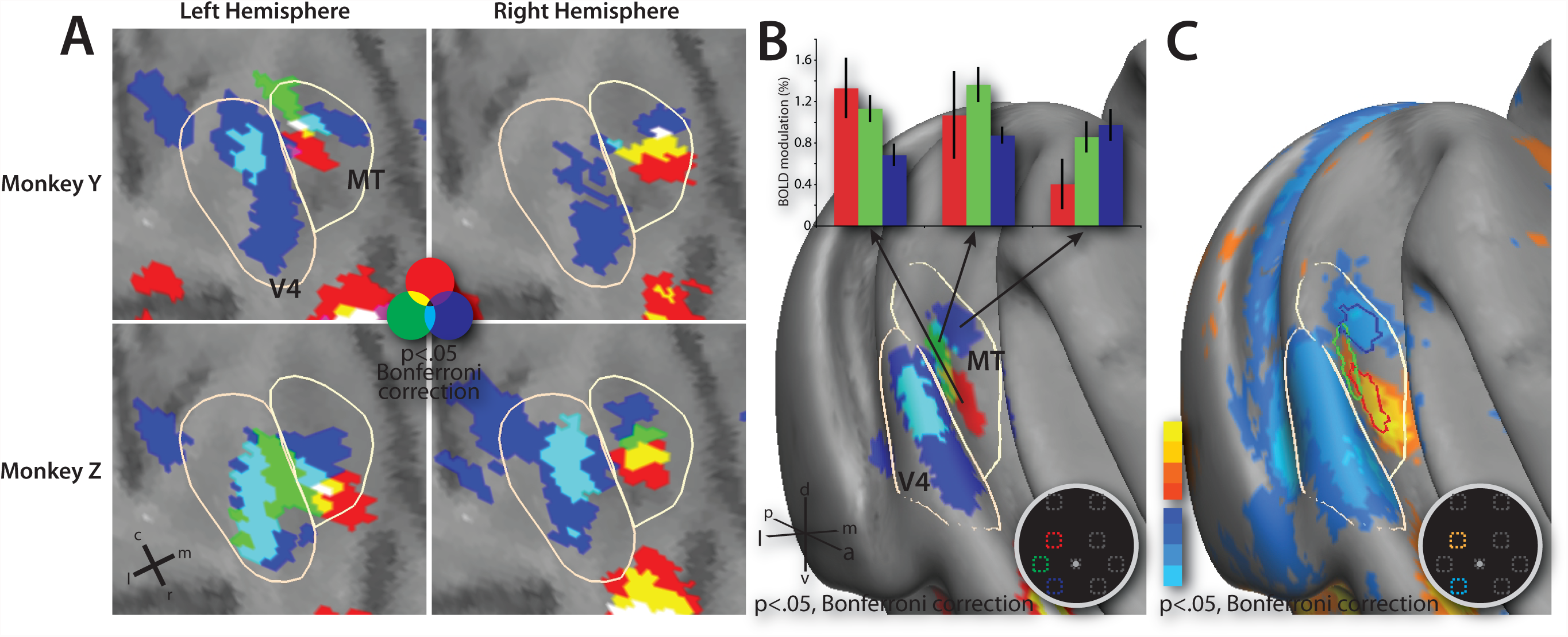
Polar angle organization of MT. **a)** Close-up of MT of the four hemispheres as projected to the macaqueF6 atlas. The beige borders outline MT and V4 as defined by the conjunction in **Figure 2**, and the dark band in the upper right is the fundus of the STS. **b)** Fixed-effects average of the MT polar-angle map project on the inflated macaqueF6 right hemisphere. Bar graph shows the magnitude of the activity evoked by the three contralateral RSVP streams for each part of the map. **c)** Upper versus lower field contrast in MT.

**Figure 6.**
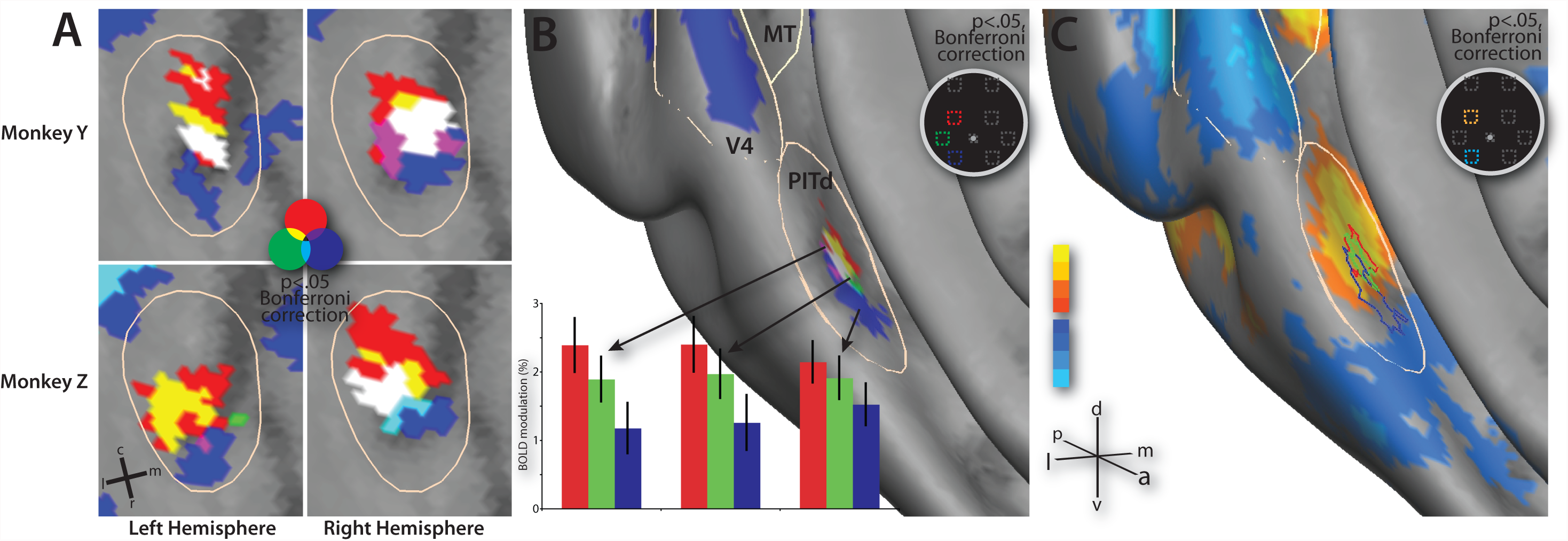
Polar angle organization of PITd. **a)** Close-up of PITd of the four hemispheres as projected to the macaqueF6 atlas. The beige borders outline PITd as defined by the conjunction in **Figure 2**, and the dark band on the right is the fundus of the STS. **b)** Fixed-effects average of the PITd polar-angle map project on the inflated macaqueF6 right hemisphere. Bar graph shows the magnitude of the activity evoked by the three contralateral RSVP streams for each part of the map. **c)** Upper versus lower field contrast in PITd.

To quantify the strength (rather than just the significance) of the topographic maps in each area, we extracted the magnitude of activity evoked by each of the six streams for each of the field representations. In LIP and MT, in addition to demonstrating that each field representation was most strongly activated by its preferred stream, the mean magnitudes revealed that the selectivity for these ROIs was strong fundus (Patel et al., 2010 Figure 3b) (see bar graph in **Figure 5.b**). In the LIP upper and lower field representations, the non-preferred stream evoked at most 1/3^rd^ as much activity as the preferred stream, and in MT the non-preferred stream evoked about ½ as much activity. The upper versus lower field maps fundus (Patel et al., 2010 Figure 3c) (**Figure 5c**) reinforced the strength of the maps in these two areas: even with a less-sensitive voxel-wise contrast, the differences in the activity evoked by the two streams were highly significant (peaks of p<.00001 Bonferroni corrected). In PITd, however, the magnitudes revealed that in all three ROIs the upper field stream evoked the most activity, despite the bias in the ROI selection and the consistency of the organization of the map across the four hemispheres (see bar graph in **Figure 6.b**). The overrepresentation of the upper field stream is corroborated by the stronger statistical scores for this stream in the individuals (see previous paragraph) and by the upper-lower field contrast (see **Figure 6.c**), which showed a strong statistical difference between the upper and lower field streams in the upper field representation but not the lower field representation.

### Periphery versus Fovea

To determine which areas contained distinct representations of the fovea and the periphery, we performed a second single-stream experiment in which the RSVP stream could appear randomly in one of three locations: at the fovea or in the periphery of the upper left or upper right field (15.6° eccentricity and 18° to the left or right of the vertical meridian). A GLM was used to create maps contrasting the activity evoked by each peripheral location versus the foveal stream for each monkey. For each hemisphere, the resulting contrast map was projected to the macaqueF6 atlas, and then thresholded to isolate the peak positive (peripheral) and negdative (foveal) areas separately for each contralateral-preferring ROI (p<.01 uncorrected). The contrast surface-maps from each hemisphere were then combined into a fixed-effects map.

Extraoccipital areas containing separate representations of the fovea and periphery included LIP, MT, PITd, FEF, and area 46p. In LIP, the peripheral representation was caudal to the foveal representation, and was slightly dorsal to and partially overlapped with the upper-field representation from the polar angle experiment Patel et al., 2010 Figures 4a and 4b)The foveal representation was rostral to the lower field representation, with little overlap; this location was unexpected as the peripheral-foveal axis in a retinotopic map is usually orthogonal to the polar-angle axis.

In MT, the peripheral representation was just medial to the upper field representation (see MT in **Figures 7.a** and **7.b**). The foveal representation was rostral and slightly lateral to the map of the periphery from the polar-angle experiment, and much of it fell in the space between MT and PITd. The peripheral representation in PITd was again slightly medial to the upper-field representation (see PITd in **Figures 7.a** and **7.b**). The PITd foveal representation in monkey Y was directly lateral to the peripheral representation, whereas in monkey Z the foveal representation was weaker and more rostral than in monkey Y. In addition, in all four hemispheres, area CITd appeared to preferentially be activated by the foveal stream over the peripheral streams (see CITd in **Figures 7.a** and **7.b**). In general, foveal representations appeared to be lateral and rostral to the peripheral representations in inferotemporal cortex.

**Figure 7.**
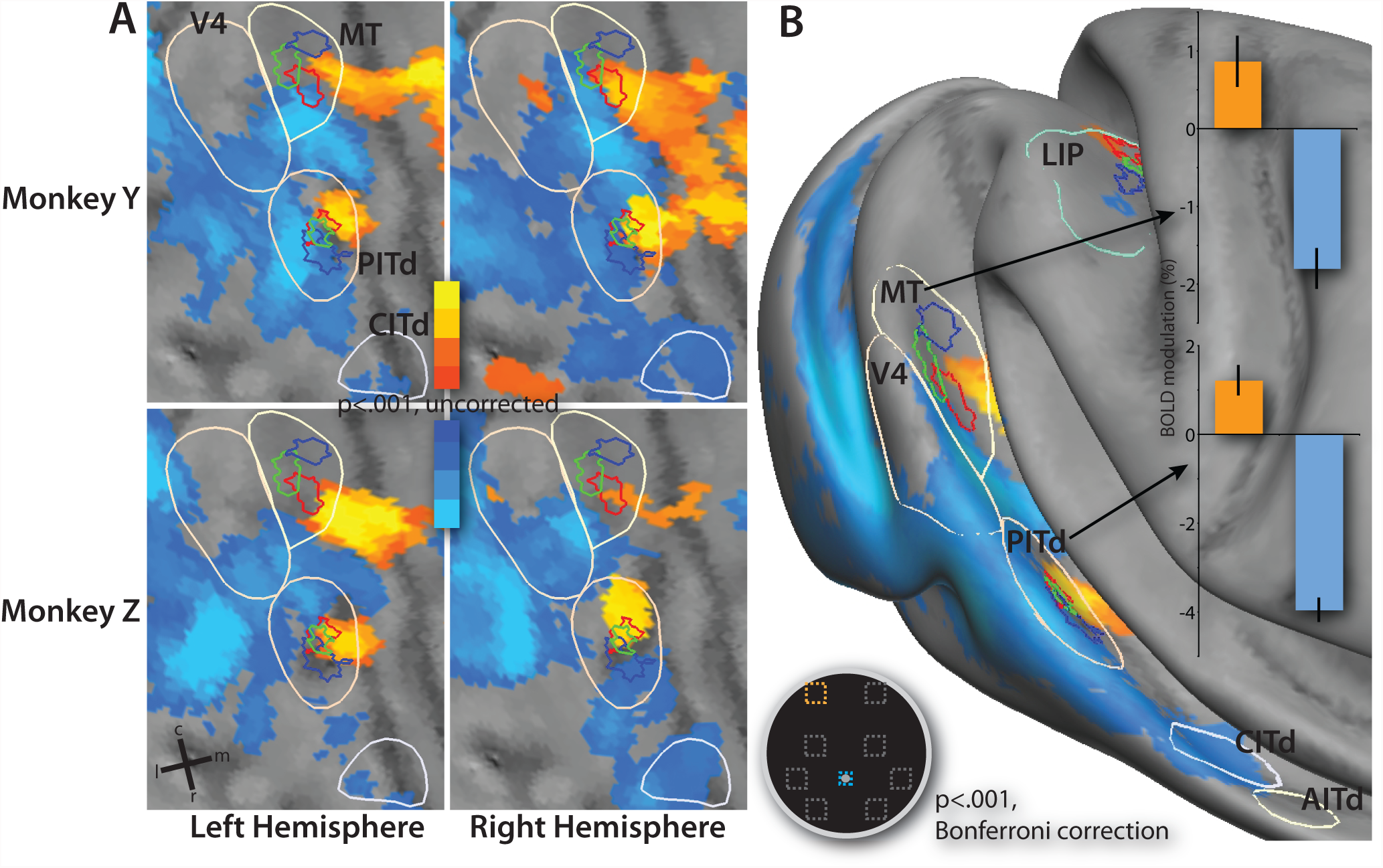
Separation of peripheral and foveal representations in MT and PITd. The red, green, and blue borders outline the upper, middle, and lower field portions of the polar angle maps, respectively, as depicted in **Figures 5** and **6**; other conventions as before. **a)** Close-up of MT, PITd, and CITd showing the peripheral (orange) and foveal (blue) representations in each of the four hemispheres. **b)** Fixed-effects average of the peripheral versus foveal stream contrast. Bar graph shows the difference in the evoked activity between the two streams for the two representations in MT and PITd.

In FEF and area 46p, the peripheral representations were dorsal to the foveal representations (see **Figures 8.a** and **8.b**). In FEF, the peripheral representation in 3 of the 4 hemispheres was slightly dorsal to the bend in the arcuate sulcus, whereas the large foveal representation was on the anterior bank of the inferior ramus of the arcuate sulcus. In the fourth hemisphere (the left hemisphere of monkey Z), there was no distinguishable representation of the periphery; the foveal representation occupied much of the anterior bank of the arcuate sulcus. In area 46p, the peripheral representation appeared on the gyral surface just posterior to the principal sulcus, and the foveal representation was ventral and anterior to this peripheral representation. In both areas, the fixed-effects map underscores the fact that while the foveal representations were strong and consistent, the peripheral representations were weaker and the location of these representations was more variable.

**Figure 8.**
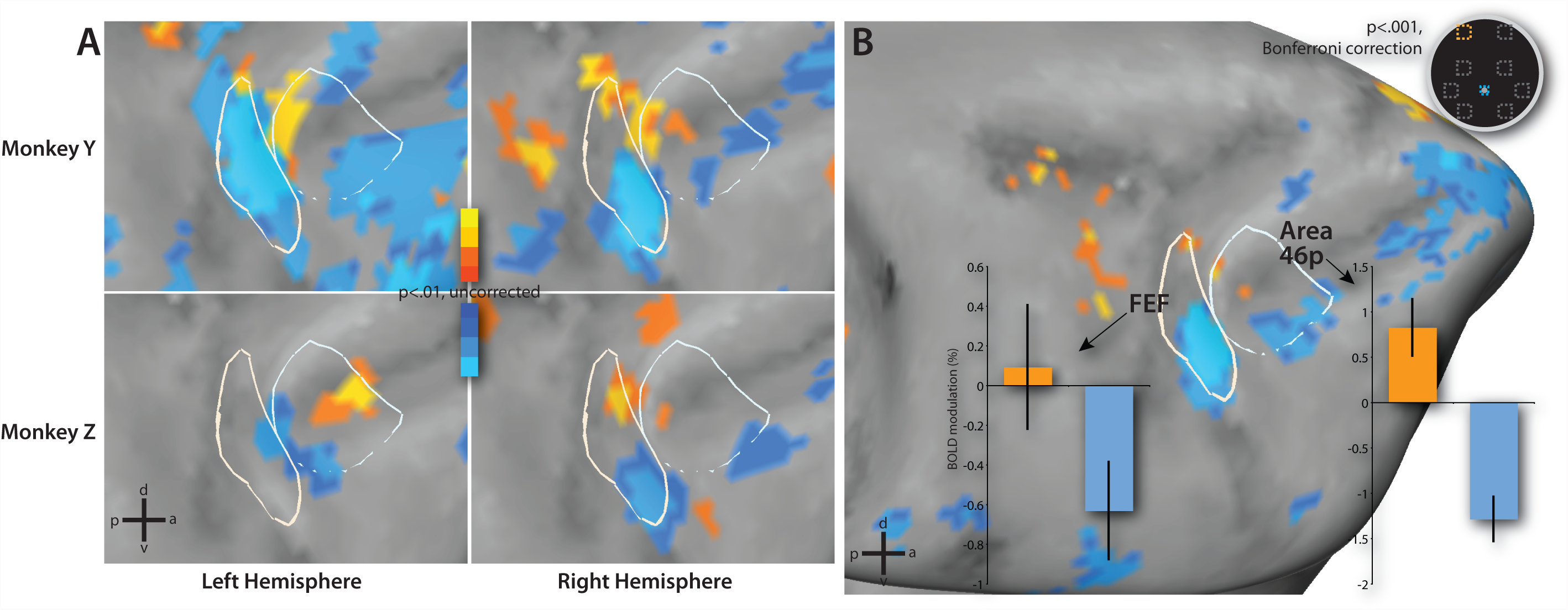
Separation of peripheral and foveal representations in FEF and area 46p. The white borders outline FEF and area 46p as defined by the conjunction map in **Figure 2.a**) Close-up of FEF and area 46p showing the peripheral (orange) and foveal (blue) representations in each of the four hemispheres. **b)** Fixed-effects average of the peripheral versus foveal stream contrast. Bar graph shows the difference in the evoked activity between the two streams for the two representations in FEF and area 46p.

To examine the strength of the peripheral and foveal biases, the fixed-effects map was used to create ROIs of the peripheral and foveal representation for each area. These ROIs were used to extract the difference in the magnitude of the activity evoked by the peripheral versus the foveal streams, with a positive difference indicating a peripheral bias and a negative difference indicating a foveal bias (see bar graphs in **Figures 7.b**, and **8.b**). The results demonstrate that for all of the areas except FEF, the bias in the foveal and peripheral representations for their preferred locations was strong and consistent.

While the fixation error was larger than desired in monkey Z, the results indicate that this did not adversely affect our results. The larger fixation error would have resulted in blurrier maps in monkey Z versus monkey Y; no such difference could be seen. In the eccentricity experiment, even if the monkey was fixating above the fixation point but within the tolerance window, the minimum distance between his eye-position and the stream was still ∼10° eccentricity. The robustness of the findings in spite of the variability in the behavior only serves to underscore the ability to generalize our findings to all macaques.

## Discussion

To investigate the retinotopic properties of higher-order visual areas, we used fMRI to measure the cortical distribution of neural activity in two macaques performing a series of demanding visuospatial attention tasks. First, we found that most visuospatial processing areas only respond to contralaterally presented stimuli; ipsilaterally presented stimuli evoked little or no activity in these areas. Second, we found that LIP, MT, and possibly PITd contained polar-angle maps of the contralateral hemifield. Third, we found that these same areas plus FEF and area 46 appear to have separate representations of the fovea and periphery. In general, we find that areas higher in the processing hierarchy tend to have coarser topography. This principle applies both in the ventral and dorsal visual stream of processing (Ungerleider and Mishkin, 1982). Moreover, when compared to previous human fMRI studies, these results indicate that there may be significant differences between macaque visual processing areas and their putative human homologues.

In the following discussion, we discuss spatial properties of visual responses separately for different areas. For each area we discuss the fMRI evidence in relation to the anatomy and physiology in macaque, then we consider functional similarities with presumptive human homologues.

### MT

MT is one of the best-characterized areas in macaque cortex, and the findings from the current study support those of many earlier electrophysiological and histological studies. Though we did not perform a specific motion localizer to isolate MT in our individual monkeys, we base our label on both the location relative to the MT boundaries in the Lewis/Van Essen and Saleem/Logothetis atlases, and the fact that in previous fMRI studies MT appeared to share a horizontal meridian with V4 (Lewis and Van Essen, 2000; Fize et al., 2003; Saleem and Logothetis, 2006). In previous histology studies, the area V4t was found to lie between MT and V4; however, given the small size of this area, it may not be distinguishable with our scanning resolution (Ungerleider et al., 2007). As expected, MT was only activated by the contralateral field streams, and represented the contralateral hemifield in retinotopic coordinates. The polar angle and eccentricity axes do not appear to be orthogonal to each other, however; the map appears to have been distorted to accommodate the shared horizontal meridian with V4, with the polar angle axis roughly at a 45° angle with respect to the eccentricity axis. This layout of the upper, middle, lower, and foveal field representations appears to match previous macaque electrophysiological and histological studies (Gattass and Gross, 1981; Van Essen et al., 1981; Desimone and Ungerleider, 1986b; Ungerleider and Desimone, 1986). Retinotopic organization and contralateral preference has been also observed in human MT; however, unlike macaque MT, human MT also appears to respond weakly to ipsilaterally-presented stimuli (Huk et al., 2002; Jack et al., 2007; Serences and Boynton, 2007). Our findings indicate that human and macaque MT appear to be similar in several aspects.

### LIP

LIP is an area often studied in the realms of visuospatial processing and oculomotor control. However, the degree to which it is topographically organized was unclear until now. Electrophysiological studies reviewed earlier found conflicting results regarding topography in LIP (Patel et al., 2010 Figure 3). In contrast, our BOLD-fMRI signals reveal a robust retinotopic map Concerning potential differences with human organization, these have been already discussed earlier, and will be only briefly reviewed here. The strongest difference concerns the location on the medial bank of putative human LIP (Sereno et al., 2001; Schluppeck et al., 2005; Silver et al., 2005; Hagler et al., 2007; Jack et al., 2007; Kastner et al., 2007; Swisher et al., 2007). The medial location indicates a potential non-linear expansion of either the lateral bank or inferior parietal lobule in humans, as suggested by interspecies warping of cortical mantles, presumably to accommodate novel verbal abilities in the human brain (Van Essen, 2003; Orban et al., 2004a).

The differences in the number of human IPS areas versus the macaque, and the potential explanations of these differences have already been discussed. In summary, the robustness of these maps, and the consistency of the polar angle and eccentricity axes of organization are not very clear in humans (Sereno et al., 2001; Schluppeck et al., 2005; Silver et al., 2005; Hagler et al., 2007; Kastner et al., 2007; Swisher et al., 2007; Orban et al., 2006; Swisher et al., 2007). Our findings support the notion of retinotopic organization in parietal cortex, but we find only one clear map in macaque IPS consistent with some of the human studies (Sereno et al., 2001; Jack et al., 2007). Another critical difference is the greater degree of ipsilateral visual response in humans (Claeys et al., 2003; Medendorp et al., 2003; Jack et al., 2007; Serences and Boynton, 2007) as compared to the macaques in this experiment. Overall, the current analysis suggests several potential differences between human and monkey visuospatial maps in IPS cortex.

### Inferotemporal Areas

While the caudal portions of the macaque STS have been extensively studied, the organization and layout of areas in the rostral portion is less clear. Various histological studies have found evidence of areal distinctions in the rostral STS, but there is some disagreement between different partitioning schemes (Desimone and Ungerleider, 1986a; Felleman and Van Essen, 1991; Lewis and Van Essen, 2000). Electrophysiological studies have not found clear functional distinctions between these areas (Desimone et al., 1984; Baylis et al., 1987; Perrett et al., 1992). Of the foci we observed in inferotemporal cortex, the most posterior fell in or near the histological boundaries of PITd, which is contained within the larger confines of the area TEO from the 1986 Ungerleider and Desimone study (Desimone and Ungerleider, 1986a). This focus overlaps with the posterior face-specific region reported in previous studies of face and object processing (Tsao et al., 2006; Op de Beeck et al., 2007). We find a complete retinotopic map contained within the boundaries of PITd; the lateral location of the fovea matches both electrophysiological recordings and more recent fMRI results (Boussaoud et al., 1991; Nelissen et al., 2006). This complete map is contained within a single focus of visually-driven activity, which further reinforces the hypothesis that this represents a single area. While the topographic map was weak in our study, the polarity of the organization was consistent in all 4 hemispheres, indicating as previous electrophysiological studies have that the cells in this area may possess larger receptive fields than the cells in earlier visual areas such as MT; large overlapping receptive fields may result in weak topographic maps when assessed by BOLD-fMRI (Boussaoud et al., 1991; Distler et al., 1993). The Boussaoud *et al*. study also found that the upper field was over-represented as compared to the lower field in this region of temporal cortex; this may explain the increased strength of the upper field representation in our study. In addition to the above findings, we find that PITd also exclusively represents the contralateral hemifield, replicating the findings of Boussaoud *et al*. and others.

Rostral to PITd were two foci of activity falling into the borders of AITd and CITd in the Felleman and Van Essen partitioning scheme; both of these foci are contained within the borders of TEc from the Ungerleider and Desimone study. Unlike the other visual processing areas, these areas did have positive responses to ipsilaterally-presented stimuli. Overall, however, these activations were less reliable than other visual processing areas—the CITd focus only appeared in 3 of the 4 hemispheres—which may have been due to their proximity to the moving jaw muscles of the monkeys. While these areas did not appear to contain retinotopic maps, CITd did appear to be more highly activated by foveal versus peripheral stimuli in the single-stream eccentricity experiment. Tsao *et al*. found that the cells in this area (labeled the “middle face-patch” in that study) are exclusively activated by faces—a highly salient stimulus—in a passive viewing task; perhaps the over-training on the identification of the 42 objects in our task has made these stimuli as salient as faces in our monkeys (Tsao et al., 2003; Tsao et al., 2006). These foci appeared to be lateral to the area LST as described by Nelissen *et al* (Nelissen et al., 2006).

In general, there appeared to be a general trend of decreasing contralateral bias the further rostral an area was located in inferotemporal cortex. This matched results of previous electrophysiology and lesion studies, which found that neurons in rostral inferotemporal areas have large receptive fields that encroach upon the ipsilateral field (Desimone et al., 1984; Desimone and Ungerleider, 1986a; Boussaoud et al., 1991; Buffalo et al., 2005). These larger receptive fields would result in ipsilateral responses in our study, which in turn would lead to a decreased contralateral index value. This trend can also be observed in human object recognition cortex, where the more rostral and higher-level area, such as the fusiform face area (FFA) (Hemond et al., 2007; Serences and Boynton, 2007), have less of a contralateral bias than more caudal regions; this may indicate some similarities between object recognition cortex in the two species.

As far as the eccentricity organization, we found that the periphery was represented medially, and the fovea was represented laterally and rostrally. The co-localization of the middle face-patch in the Tsao *et al*. study with the foveal-preferring AITd/CITd foci in our study may be consistent with the observation in humans that face-processing regions have a foveal bias (Hasson et al., 2002). However, in humans the foveal and peripheral representations in object recognition areas are continuous with the earlier visual areas (Hasson et al., 2002; Levy et al., 2004; Hansen et al., 2007). In our data, while the foveal representations in inferotemporal cortex appear to be continuous with the fovea in earlier visual areas, the peripheral representations are situated medial to the fovea, rather than lateral to the fovea as predicted by the human precedent.

Overall, these results give a mixed picture of the homologous relationship of object recognition areas in the two species. The similarities in contralateral bias may indicate that the cells underlying the processing of one part of the visual field are similarly organized between the two species. However, the differences in the foveal-peripheral organization may indicate radical differences in the way these difference patches of cortex are connected, which in turn may point to differences in the mechanisms underlying visual processing.

### Prefrontal Cortex

In prefrontal cortex, there were two main foci of activity in all four hemispheres. In the arcuate sulcus, the boundaries of the evoked BOLD signal in our study matched those of FEF from previous studies (Bruce et al., 1985; Felleman and Van Essen, 1991; Saleem and Logothetis, 2006). Compared to previous fMRI studies, however, the boundaries in our conjunction map are more restrictive (Koyama et al., 2004; Baker et al., 2006). This difference holds true even for the main effect of visual stimulation (activity evoked by the streams in the two-stream paradigm), and did not appear to be an effect of the threshold level. The focal activation of FEF may reflect the specificity of visual responses, while previous studies combined visual and oculomotor activity. Furthermore, this is also the first study to explicitly separate effects of reward from the effects of visual stimulation. The second focus of activity corresponded to the ventral portion of area 46p, which matched with the location of electrophysiology studies of working memory and object discrimination performed by Miller *et al*. and *WIlson et al*. (Wilson et al., 1993; Miller et al., 1996).

The exclusive representation in both areas of the contralateral field matches previous macaque electrophysiological (and in the case of FEF, electrical stimulation) studies of both areas (Bruce et al., 1985; Funahashi et al., 1989; Chafee and Goldman-Rakic, 1998; Sommer and Wurtz, 2000). Previous studies have found a clear eccentricity map in FEF but differ as to whether there is an accompanying polar angle map (Bruce et al., 1985; Sommer and Wurtz, 2000); our results support the former but not the latter claim for both FEF and area 46p. However, the peripheral representation in both areas is weak in our study, especially in FEF. This weakness may be due to the geometry of the cortical surface in this region due to the arrangement of the arcuate and principal sulci: complex folding within the space of a couple voxels may obscure and confuse underlying activations. This may hold true for the polar angle map, so the lack of a polar angle map in our study in either of these regions must be taken with a note of caution. In general the foveal representation appeared to be larger in surface area than the peripheral representation in both areas, possibly pointing to an overrepresentation of the fovea. An oculomotor working memory study by Sawaguchi and Iba found a retinotopic map on the dorsal bank of the principal sulcus; we did not find any evidence of a map in this area (Sawaguchi and Iba, 2001).

The locations of human prefrontal cortex areas involved in visuospatial processing have been well characterized. They include a putative homologue of macaque area 46 on the middle frontal gyrus, and the putative FEF homologue at the junction of the superior frontal and precentral sulci (Corbetta et al., 1998; Petrides and Pandya, 1999; Tehovnik et al., 2000; Beauchamp et al., 2001; Orban et al., 2004b). Both of these human areas demonstrate a strong preference for the contralateral field, though in contrast with their macaque homologues, they also both respond to ipsilateral stimuli (Medendorp et al., 2003; Jack et al., 2007; Serences and Boynton, 2007). Recent studies have claimed to have also found retinotopic maps in both of these areas, with both polar-angle and eccentricity axes of organization (Hagler and Sereno, 2006; Hagler et al., 2007; Kastner et al., 2007). Work from our own lab, however, suggests that these maps are weak if they even exist at all (Jack et al., 2007). In addition, work from several labs suggest that the human FEF homologue contains two separate functional regions, though it is unclear what if any functional distinctions exist between these two locations (Corbetta et al., 1998; Beauchamp et al., 2001; Jack et al., 2007). If all of these characteristics of the human prefrontal areas prove to be true, then it may indicate that substantial differences exist in prefrontal cortex between the two species.

### Conclusions

We found that areas at higher levels of the visual hierarchy show coarser topographic visual responses for stimuli in the contralateral visual field, and even less contralateral specificity: this is true in both the ventral stream (PITd-CITd/AITd-46p) and the dorsal stream (MT-LIP-FEF). This decrement in spatial specificity arises from the overlap of increasingly larger receptive fields and/or from increasing mixing of cells representing different parts of the visual field. It has been proposed that topographic maps underlie a very specific computational and energetic principle, wiring optimization, according to which neurons that perform similar local operations are heavily connected and therefore packed in adjacent portions of cortex. Conversely, neurons in areas that perform more global operations, such as integrating information from different locations in the visual field or associating flexibly a stimulus to different responses, have fewer local connections but more connections with other areas. This principle explains why higher-order areas that are important for integrating information within a sensory system or for associating stimuli to response may not be strongly topographic (Chklovskii and Koulakov, 2004).

The results of this study also bear on three points about the evolution of the cortex between macaques and humans. First, the most striking inter-species difference is the relative location of different higher order visual areas in relation to the major sulci and gyri. In temporal cortex, inferotemporal areas are laterally located in monkeys while they are more ventral and medial in humans (Denys et al., 2004; Orban et al., 2004b). In parietal cortex, LIP is on the lateral bank while the presumptive human homologue is on the medial bank (Sereno et al., 2001; Denys et al., 2004; Orban et al., 2004b). In prefrontal cortex areas, FEF and 46 are adjacent in monkeys while in humans are separated so that 46 is on the middle frontal gyrus while FEF is more posteriorly located at the intersection of precentral and superior frontal sulcus (Petrides and Pandya, 1999; Tehovnik et al., 2000). These results indicate the non-linear expansion of lateral temporal cortex (between A1 and the ventral stream areas), temporoparietal cortex (between MT and LIP), and prefrontal cortex (between FEF and area 46) in humans likely associated to the development of verbal abilities, working memory, and self-referential processing (Van Essen, 2003).

Second, unlike higher-level human visual processing areas, macaque areas do not respond to the presentation of ipsilateral stimuli (with the exception of rostral temporal cortex). The ipsilateral responses in humans do not appear to form a topographic map, and may represent signals for suppressing irrelevant stimuli from unattended parts of the visual field or for integrating more flexibly information across visual fields. This may indicate either a difference in attentional control or cognitive strategies. It may also indicate the different degree of practice with the task since in humans ipsilateral responses are measured after hundreds of trials while monkeys performed many thousands of trials before being scanned (Bichot et al., 1996).

Third, there is some evidence for an increase in the number of visuospatial areas within human IPS and near/at FEF. The evidence in humans, however, will need further confirmation and more direct comparisons with monkeys using identical tasks. If true, this interspecies difference may also indicate a difference in the mechanisms underlying basic visuospatial tasks. Differences in these mechanisms between the humans and macaques are partly supported by recently described behavioral differences in the execution of similar cognitive operations (Stoet and Snyder, 2007). While we are still at a very recent early stage of development of a comparative approach of neurocognitive systems in humans and monkeys, in our view these results suggest that caution must be exerted in applying macaque data to human models of cognition.

## References

Baker JT, Patel GH, Corbetta M, Snyder LH (2006) Distribution of Activity Across the Monkey Cerebral Cortical Surface, Thalamus and Midbrain during Rapid, Visually Guided Saccades. Cereb Cortex 16:447–459.

Bakola S, Gregoriou GG, Moschovakis AK, Savaki HE (2006) Functional imaging of the intraparietal cortex during saccades to visual and memorized targets. Neuroimage 31:1637–1649.

Baylis GC, Rolls ET, Leonard CM (1987) Functional subdivisions of the temporal lobe neocortex. J Neurosci 7:330–342.

Beauchamp MS, Petit L, Ellmore TM, Ingeholm J, Haxby JV (2001) A parametric fMRI study of overt and covert shifts of visuospatial attention. Neuroimage 14:310–321.

Ben Hamed S, Duhamel JR, Bremmer F, Graf W (2001) Representation of the visual field in the lateral intraparietal area of macaque monkeys: a quantitative receptive field analysis. Exp Brain Res 140:127–144.

Bichot NP, Schall JD, Thompson KG (1996) Visual feature selectivity in frontal eye fields induced by experience in mature macaques. Nature 381:697–699.

Blatt GJ, Andersen RA, Stoner GR (1990) Visual receptive field organization and cortico-cortical connections of the lateral intraparietal area (area LIP) in the macaque. Journal of Comparative Neurology 299:421–445.

Boussaoud D, Desimone R, Ungerleider LG (1991) Visual topography of area TEO in the macaque. J Comp Neurol 306:554–575.

Boynton GM, Engel SA, Glover GH, Heeger DJ (1996) Linear systems analysis of functional magnetic resonance imaging in human V1. Journal of Neuroscience 16:4207–4221.

Brewer AA, Press WA, Logothetis NK, Wandell BA (2002) Visual areas in macaque cortex measured using functional magnetic resonance imaging. J Neurosci 22:10416–10426.

Bruce CJ, Goldberg ME, G.B. S (1985) Primate frontal eye fields. II. Physiological and anatomical correlates of electrically evoked eye movements. Journal of Neurophysiology 54:714–734.

Buffalo EA, Bertini G, Ungerleider LG, Desimone R (2005) Impaired filtering of distracter stimuli by TE neurons following V4 and TEO lesions in macaques. Cereb Cortex 15:141–151.

Chafee MV, Goldman-Rakic PS (1998) Matching patterns of activity in primate prefrontal area 8a and parietal area 7ip neurons during a spatial working memory task. J Neurophysiol 79:2919–2940.

Chklovskii DB, Koulakov AA (2004) Maps in the brain: what can we learn from them? Annu Rev Neurosci 27:369–392.

Claeys KG, Lindsey DT, De Schutter E, Orban GA (2003) A higher order motion region in human inferior parietal lobule: evidence from fMRI. Neuron 40:631–642.

Corbetta M, Akbudak E, Conturo TE, Snyder AZ, Ollinger JM, Drury HA, Linenweber MR, Petersen SE, Raichle ME, Van Essen DC, Shulman GL (1998) A common network of functional areas for attention and eye movements. Neuron 21:761–773.

Denys K, Vanduffel W, Fize D, Nelissen K, Peuskens H, Van Essen D, Orban GA (2004) The processing of visual shape in the cerebral cortex of human and nonhuman primates: a functional magnetic resonance imaging study. J Neurosci 24:2551–2565.

Desimone R, Ungerleider LG (1986a) Multiple visual areas in the caudal superior temporal sulcus of the macaque. Journal of Comparative Neurology 248:164–189.

Desimone R, Ungerleider LG (1986b) Multiple visual areas in the caudal superior temporal sulcus of the macaque. J Comp Neurol 248:164–189.

Desimone R, Albright TD, Gross CG, Bruce C (1984) Stimulus-selective properties of inferior temporal neurons in the macaque. J Neurosci 4:2051–2062.

DeYoe E, Carman G, Bandettini, Glickman S, Wieser J, Cox R, Miller D, Neitz J (1996) Mapping striate and extrastriate visual areas in human cerebral cortex. Proceedings of the National Academy of Sciences of the United States of America 93:2382–2386.

Distler C, Boussaoud D, Desimone R, Ungerleider LG (1993) Cortical connections of inferior temporal area TEO in macaque monkeys. J Comp Neurol 334:125–150.

Felleman DJ, Van Essen DC (1991) Distributed hierarchical processing in the primate cerebral cortex. Cerebral Cortex 1:1–47.

Fize D, Vanduffel W, Nelissen K, Denys K, Chef d’Hotel C, Faugeras O, Orban GA (2003) The retinotopic organization of primate dorsal V4 and surrounding areas: A functional magnetic resonance imaging study in awake monkeys. J Neurosci 23:7395–7406.

Funahashi S, Bruce CJ, Goldman-Rakic PS (1989) Mnemonic coding of visual space in the monkey’s dorsolateral prefrontal cortex. Journal of Neurophysiology 61:331–349.

Gattass R, Gross CG (1981) Visual topography of striate projection zone (MT) in posterior superior temporal sulcus of the macaque. J Neurophysiol 46:621–638.

Hagler DJ, Jr., Sereno MI (2006) Spatial maps in frontal and prefrontal cortex. Neuroimage 29:567–577.

Hagler DJ, Jr., Riecke L, Sereno MI (2007) Parietal and superior frontal visuospatial maps activated by pointing and saccades. Neuroimage 35:1562–1577.

Hansen KA, Kay KN, Gallant JL (2007) Topographic organization in and near human visual area V4. J Neurosci 27:11896–11911.

Hasson U, Levy I, Behrmann M, Hendler T, Malach R (2002) Eccentricity bias as an organizing principle for human high-order object areas. Neuron 34:479–490.

Hemond CC, Kanwisher NG, Op de Beeck HP (2007) A preference for contralateral stimuli in human object- and face-selective cortex. PLoS ONE 2:e574.

Huk AC, Dougherty RF, Heeger DJ (2002) Retinotopy and functional subdivision of human areas MT and MST. J Neurosci 22:7195–7205.

Jack AI, Patel GH, Astafiev SV, Snyder AZ, Akbudak E, Shulman GL, Corbetta M (2007) Changing human visual field organization from early visual to extra-occipital cortex. PLoS ONE 2:e452.

Kastner S, DeSimone K, Konen CS, Szczepanski SM, Weiner KS, Schneider KA (2007) Topographic maps in human frontal cortex revealed in memory-guided saccade and spatial working-memory tasks. J Neurophysiol 97:3494–3507.

Koyama M, Hasegawa I, Osada T, Adachi Y, Nakahara K, Miyashita Y (2004) Functional magnetic resonance imaging of macaque monkeys performing visually guided saccade tasks: comparison of cortical eye fields with humans. Neuron 41:795–807.

Levy I, Hasson U, Harel M, Malach R (2004) Functional analysis of the periphery effect in human building related areas. Hum Brain Mapp 22:15–26.

Lewis JW, Van Essen DC (2000) Mapping of architectonic subdivisions in the macaque monkey, with emphasis on parieto-occipital cortex. Journal of Comparative Neurology 428:79–111.

Logothetis NK, Wandell BA (2004) Interpreting the BOLD signal. Annu Rev Physiol 66:735–769.

Medendorp WP, Goltz HC, Vilis T, Crawford JD (2003) Gaze-centered updating of visual space in human parietal cortex. J Neurosci 23:6209–6214.

Miller EK, Erickson CA, Desimone R (1996) Neural mechanisms of visual working memory in prefrontal cortex of the macaque. Journal of Neuroscience 16:5154–5167.

Nelissen K, Vanduffel W, Orban GA (2006) Charting the lower superior temporal region, a new motion-sensitive region in monkey superior temporal sulcus. J Neurosci 26:5929–5947.

Op de Beeck HP, Deutsch JA, Vanduffel W, Kanwisher NG, Dicarlo JJ (2007) A Stable Topography of Selectivity for Unfamiliar Shape Classes in Monkey Inferior Temporal Cortex. Cereb Cortex.

Orban GA, Van Essen D, Vanduffel W (2004a) Comparative mapping of higher visual areas in monkeys and humans. Trends Cogn Sci 8:315–324.

Orban GA, Van Essen D, Vanduffel W (2004b) Comparative mapping of higher visual areas in monkeys and humans. Trends Cogn Sci 8:315–324.

Orban GA, Claeys K, Nelissen K, Smans R, Sunaert S, Todd JT, Wardak C, Durand JB, Vanduffel W (2006) Mapping the parietal cortex of human and non-human primates. Neuropsychologia 44:2647–2667.

Patel GH (2010) Topographic organization of the macaque area LIP. Proc Natl Acad Sci USA:107(10)4728–33.

Perrett DI, Hietanen JK, Oram MW, Benson PJ (1992) Organization and functions of cells responsive to faces in the temporal cortex. Philos Trans R Soc Lond B Biol Sci 335:23–30.

Petrides M, Pandya DN (1999) Dorsolateral prefrontal cortex: comparative cytoarchitectonic analysis in the human and the macaque brain and corticocortical connection patterns. Eur J Neurosci 11:1011–1036.

Platt ML, Glimcher PW (1998) Response fields of intraparietal neurons quantified with multiple saccadic targets. Exp Brain Res 121:65–75.

Saleem KS, Logothetis NK (2006) A Combined MRI and Histology Atlas of the Rhesus Monkey Brain: Academic Press.

Sawaguchi T, Iba M (2001) Prefrontal cortical representation of visuospatial working memory in monkeys examined by local inactivation with muscimol. J Neurophysiol 86:2041–2053.

Schluppeck D, Glimcher P, Heeger DJ (2005) Topographic organization for delayed saccades in human posterior parietal cortex. J Neurophysiol 94:1372–1384.

Serences JT, Boynton GM (2007) Feature-based attentional modulations in the absence of direct visual stimulation. Neuron 55:301–312.

Sereno MI, Tootell RB (2005) From monkeys to humans: what do we now know about brain homologies? Curr Opin Neurobiol 15:135–144.

Sereno MI, Pitzalis S, Martinez A (2001) Mapping of contralateral space in retinotopic coordinates by a parietal cortical area in humans. Science 294:1350–1354.

Sereno MI, Dale AM, Reppas JB, Kwong KK, Belliveau JW, Brady TJ, Rosen BR, Tootell RBH (1995) Borders of multiple visual areas in humans revealed by functional magnetic resonance imaging. Science 268:889–893.

Silver MA, Ress D, Heeger DJ (2005) Topographic maps of visual spatial attention in human parietal cortex. J Neurophysiol 94:1358–1371.

Snyder AZ (1995) Difference image vs. ratio image error function forms in PET-PET realignment. In: Quantification of Brain Function Using PET (Myer R, Cunningham VJ, Bailey DL, Jones T, eds), pp 131–137. San Diego, CA: Academic Press.

Sommer MA, Wurtz RH (2000) Composition and topographic organization of signals sent from the frontal eye field to the superior colliculus. J Neurophysiol 83:1979–2001.

Stoet G, Snyder LH (2007) Extensive practice does not eliminate human switch costs. Cogn Affect Behav Neurosci 7:192–197.

Swisher JD, Halko MA, Merabet LB, McMains SA, Somers DC (2007) Visual topography of human intraparietal sulcus. J Neurosci 27:5326–5337.

Tehovnik EJ, Sommer MA, Chou IH, Slocum WM, Schiller PH (2000) Eye fields in the frontal lobes of primates. Brain Res Brain Res Rev 32:413–448.

Tootell RB, Tsao D, Vanduffel W (2003) Neuroimaging weighs in: humans meet macaques in ‘Primate’ visual cortex. J Neurosci 23:3981–3989.

Tsao DY, Freiwald WA, Tootell RB, Livingstone MS (2006) A cortical region consisting entirely of face-selective cells. Science 311:670–674.

Tsao DY, Freiwald WA, Knutsen TA, Mandeville JB, Tootell RB (2003) Faces and objects in macaque cerebral cortex. Nat Neurosci 6:989–995.

Ungerleider LG, Mishkin M (1982) Two cortical visual systems. In: Analysis of Visual Behavior (Ingle DJ, Goodale MA, Mansfield RJW, eds), pp 549–580. Cambridge, Mass: MIT Press.

Ungerleider LG, Desimone R (1986) Cortical connections of visual area MT in the macaque. Journal of Comparative Neurology 248:190–222.

Ungerleider LG, Galkin TW, Desimone R, Gattass R (2007) Cortical Connections of Area V4 in the Macaque. Cereb Cortex.

Van Essen D (2002) Windows on the brain: the emerging role of atlases and databases in neuroscience. Curr Opin Neurobiol 12:574–579.

Van Essen DC (2003) Organization of visual areas in Macaque and human cerebral cortex. In: The Visual Neurosciences (Werner LCJS, ed): MIT Press.

Van Essen DC, Maunsell JHR, Bixby JL (1981) The middle temporal visual area in the macaque: myeloarchitecture, connections, functional properties and topographic organization. Journal of Comparative Neurology 199:293–326.

Viswanathan A, Freeman RD (2007) Neurometabolic coupling in cerebral cortex reflects synaptic more than spiking activity. Nat Neurosci 10:1308–1312.

Wandell BA, Dumoulin SO, Brewer AA (2007) Visual field maps in human cortex. Neuron 56:366–383.

Wilson FAW, O’Scalaidhe SP, Goldman-Rakic PS (1993) Dissociation of object and spatial processing domains in primate prefrontal cortex. Science 260:1955–1958.

